# Posttranscriptional regulation of cell wall integrity in budding yeast

**DOI:** 10.1101/2022.09.30.510326

**Authors:** Stefan Bresson, Vadim Shchepachev, David Tollervey

## Abstract

The fungal cell wall provides protection and structure, and is an important target for antifungal compounds. A MAP kinase cascade termed the cell wall integrity (CWI) pathway regulates transcriptional responses to cell wall damage. Here we describe a posttranscriptional pathway that plays an important complementary role. We discovered that the RNA-binding proteins (RBPs) Mrn1 and Nab6 specifically target the 3’ UTRs of a largely overlapping set of cell wall-related mRNAs. These mRNAs are downregulated in the absence of Nab6, indicating a function in target mRNA stabilization. Nab6 acts in parallel to CWI signaling to maintain appropriate expression of cell wall genes during stress. Cells lacking both pathways are hypersensitive to antifungal compounds targeting the cell wall. Deletion of *MRN1* partially alleviates growth defects associated with *Δnab6* and Mrn1 has an opposing function in mRNA destabilization. Our results uncover a novel posttranscriptional pathway which mediates cellular resistance to antifungal compounds.

## INTRODUCTION

Most organisms growing in the wild will frequently encounter a range of environmental stresses. This is particularly the case for budding yeasts that grow on the surface of plants, including fruits, grains and bark. The fungal cell wall provides the primary physical barrier to the external environment and confers substantial protection against environmental insults. In *Saccharomyces cerevisiae*, the cell wall is comprised of an inner layer of branched β-1,3 glucan, β-1,6 glucan, and chitin, and a covalently attached outer layer of mannoproteins (Gow *et al*, 2017). The inner layer provides rigidity and structural support, while the outer layer allows the cell to interact with the external environment. Most components of the cell wall are unique to fungi, and thus form common substrates for pathogen-associated pattern recognition receptors (PRRs) in plant and animal immune systems. For the same reason, the cell wall is an attractive target for antifungal drugs. The echinocandin family of compounds, including caspofungin, inhibits β-1,3 glucan synthase (Szymanski *et al*, 2022). Other antifungal compounds such as Congo Red and Calcofluor White seem to disrupt cell wall structure by binding chitin and β-glucan polymers (Elorza *et al*, 1983; Roncero & Duran, 1985).

The major signaling pathway regulating cell wall growth and homeostasis is the cell wall integrity (CWI) pathway (Sanz *et al*, 2022). Damage to the cell wall is detected by several sensor proteins (Wsc1-3, Mid2, and Mtl1) located in the plasma membrane. These stimulate the GEF/GTPase pair Rom2 and Rho1, which in turn activate the Pkc1 kinase. Pkc1 initiates a mitogen-activated protein kinase (MAPK) cascade consisting of Bck1, Mkk1/Mkk2, and Slt2. Ultimately, Slt2 launches a transcriptional response via the transcription factor Rlm1 and the chromatin modifiers SWI/SNF and SAGA (Garcia *et al*, 2004; Sanz *et al*, 2012). Slt2 also regulates RNA polymerase II directly, by phosphorylating Tyr1 residues in the C-terminal domain (Yurko *et al*, 2017). CWI signaling is supplemented by the high-osmolarity glycerol (HOG) pathway, which responds to a subset of cell wall stresses, most notably zymolyase treatment (Bermejo *et al*, 2008; Garcia *et al*, 2015).

Posttranscriptional pathways may provide an additional layer of regulation in cell wall biosynthesis and/or integrity (Hall & Wallace, 2022). A previous RNA-immunoprecipitation and microarray screen identified several RNA binding proteins (RBPs), including Bfr1, Hek2, Mrn1, Nab6, Pub1, Scp160, and Ssd1, that preferentially associate with cell wall-related mRNAs (Hogan *et al*, 2008). The best characterized of these proteins is Ssd1, which represses translation initiation of specific cell wall mRNAs, probably via its interactions with the 5’ UTRs of these transcripts (Bayne *et al*, 2022; Hu *et al*, 2018; Jansen *et al*, 2009). Notably, the *ssd1Δ* knockout confers sensitivity to compounds targeting the cell wall (Bayne *et al.*, 2022; Jansen *et al.*, 2009; Lee *et al*, 2014), and synergistically impairs growth when combined with mutations in the CWI pathway (Costanzo *et al*, 2016). Other RBPs targeting cell wall mRNAs have not yet been investigated in detail, but may have similar roles in regulating cell wall growth and integrity

We previously assessed changes in global RNA-protein interactions following a variety of stresses using total RNA-associated proteome purification (TRAPP) (Bresson *et al*, 2020; Shchepachev *et al*, 2019). Among the proteins showing stress-induced changes in RNA binding were the putative cell-wall regulators Mrn1 and Nab6. To better understand their roles, we characterized the RNA targets for Mrn1 and Nab6. We report that both proteins specifically target the 3’ UTRs of a largely overlapping group of cell wall-related mRNAs, but have antagonistic effects on gene expression.

## RESULTS

### Mrn1 and Nab6 are related proteins with similar domain architectures

Nab6 and Mrn1 were previously predicted to contain two and four RNA-recognition motifs (RRMs), respectively (Fig. 1A). By analyzing the Alphafold prediction for Nab6 (Jumper *et al*, 2021; Varadi *et al*, 2022), we identified two additional RRMs (designated ‘putative’ and shown in blue and white stripes) which show robust structural homology to a conventional RRM (Fig. 1A, S1A-D). A BLAST search for Nab6 homologs across the *S. cerevisiae* genome returned Mrn1 as the top hit, suggesting a close evolutionary relationship between the two proteins. Further analysis revealed that Nab6 and Mrn1 share a region of homology encompassing RRM3 and RRM4 in each protein (Fig. S1E-F). Taken together, these observations suggest both proteins likely function through direct RNA binding, and possibly share related functions and/or RNA targets. Among fungi, Mrn1 is highly conserved whereas Nab6 shows lower conservation, but is clearly present in pathogens, including *Candida* sp. Both proteins have only limited homology to human RNA binding proteins.

**Figure 1.**
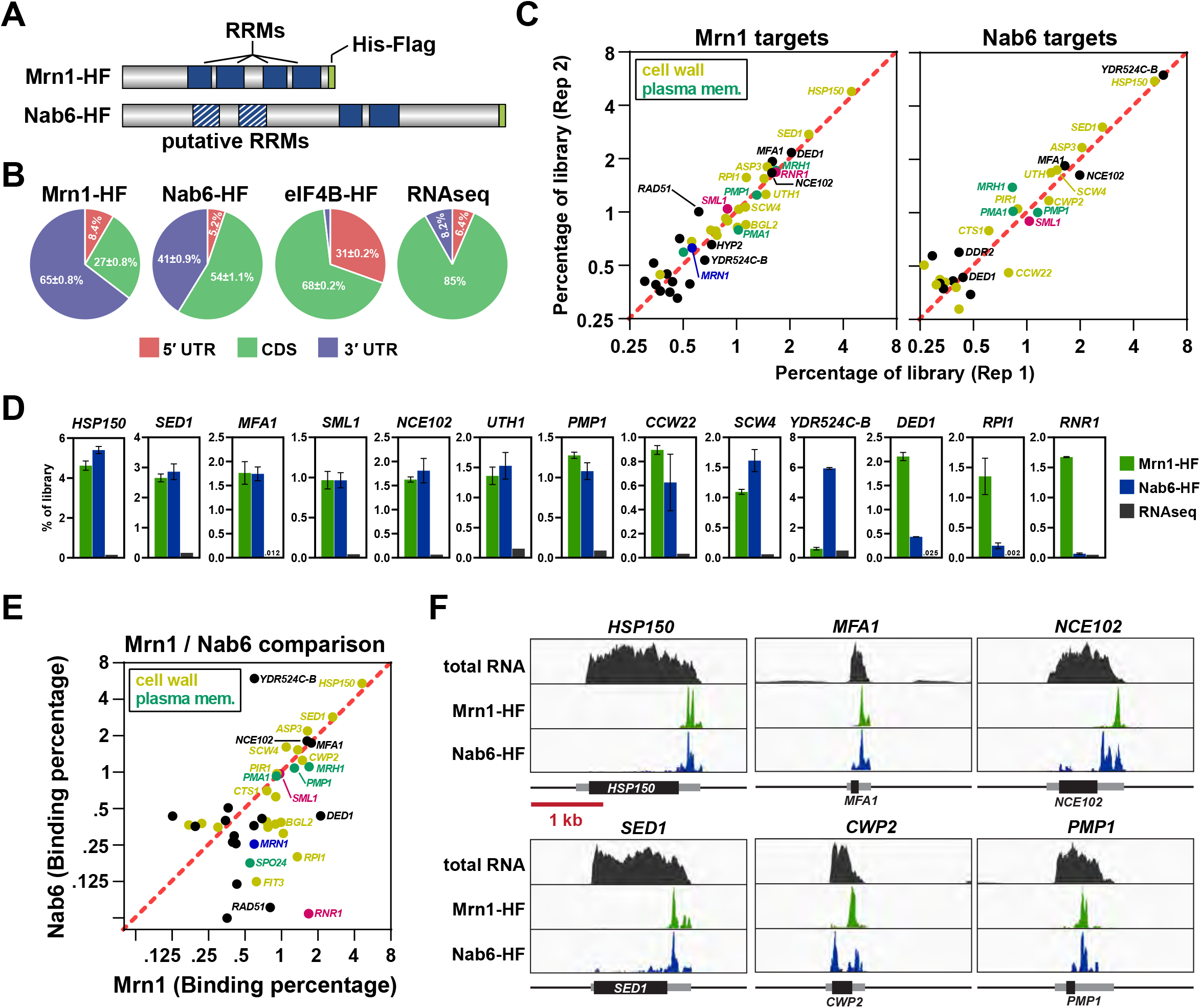
Transcriptome-wide identification of Mrn1 and Nab6 targets. (A) Domain architecture of Mrn1 and Nab6. Two putative RRMs identified using Alphafold are shown in blue and white stripes. Each protein was tagged with His8-Flag at the C-terminus. (B) Distribution of Mrn1, Nab6, and eIF4B binding to the 5’ UTR, CDS, and 3’ UTR regions of target RNAs. RNAseq data is included as a control. (C) Scatter plots showing the top mRNA targets of Mrn1 *(left)* and Nab6 *(right).* Each plot includes all genes comprising at least 0.35% of mRNA-mapped reads. See Table S1 for a complete list of target transcripts. (D) Bar graphs showing the percentage of reads in each library which map to selected transcripts. Error bars show standard deviation of the mean (n=2) (E) Comparison of the top targets of Mrn1 and Nab6. (F) Binding of Mrn1 and Nab6 across shared target transcripts. Each track is normalized to total library size using reads per million. RNAseq reads are included as a control. Each box represents a 3 kb window; a scale bar is shown at the bottom. The open reading frames (ORFs) are shown as black boxes, with UTRs as flanking gray boxes. Each transcript is oriented 5’ to 3’.

### Mrn1 and Nab6 target the 3’ UTRs of cell wall-related mRNAs

To identify the RNA targets of Mrn1 and Nab6, we mapped their RNA binding sites using CRAC (Granneman *et al*, 2009). Mrn1 and Nab6 were separately expressed as C-terminal HF-tagged (His8-Ala4-Flag) fusion proteins under the control of their endogenous promoters (Fig. 1A). The resulting strains were grown to exponential phase and UV-irradiated to covalently fix direct protein-RNA contacts. Subsequently, RNAs associated with Mrn1 or Nab6 were isolated using tandem affinity purification, treated with RNase to generate protein-protected RNA footprints, and analyzed by high-throughput sequencing.

We first examined the distribution of sequencing reads across 5’ UTR, coding region, and 3’ UTR sequences (Fig. 1B). Mrn1 and Nab6 preferentially bound 3’ UTRs; these formed 65% and 41 % of CRAC reads, respectively, compared to just 8% of total RNA sequencing reads (RNAseq). As a control, we analyzed CRAC reads derived from the translation initiation factor eIF4B, which was also HF-tagged (Bresson *et al.*, 2020). As expected, eIF4B was strongly enriched at 5’ UTRs, and showed negligible binding to 3’ UTRs (Fig. 1B).

Mrn1 and Nab6 reproducibly bound a relatively small set of transcripts (Figs. 1C-E and Table S1), with substantial overlap in their targets. Both proteins bound a core set of mRNAs encoding cell wall components (highlighted in yellow), including *HSP150, SED1, CWP2, ASP3*, and *SCW4* among others. Mrn1 additionally targeted *RPI1*, encoding a transcription factor for cell wall proteins (Chin *et al*, 2012). The most enriched Nab6 target was *YDR524C-B.* Although designated as an uncharacterized gene, YDR524C-B shows strong genetic interactions with various cell wall factors (Fig. S2A) (Costanzo *et al.*, 2016). In addition to cell wall-related mRNAs, several other targets were identified (Figs. 1C-E), including transcripts for plasma membrane proteins (highlighted in green). Mrn1 also targeted *RNR1*, encoding ribonucleotide reductase, and both Mrn1 and Nab6 bound *SML1*, encoding an Rnr1 inhibitor (both highlighted in pink). Another prominent target for both Mrn1 and Nab6 was *MFA1*, encoding a secreted mating pheromone. Mrn1 also bound its own mRNA, but within the 5’ UTR, indicative of possible autoregulation (Fig. S2B).

RBPs often interact promiscuously with a wide range of transcripts. However, comparing representation in CRAC with mRNA abundance measured by RNAseq indicated that Mrn1 and Nab6 show surprisingly high specificity for their mRNA targets (Fig. 1D and Table S1). For example, reads mapping to the *HSP150* mRNA comprised ~5% of the Mrn1 and Nab6 CRAC libraries, but only 0.15% of all mRNA reads. Moreover, cell wall mRNAs, as a class, were highly overrepresented in both CRAC datasets (Fig. S2C).

Finally, we examined the distribution of Mrn1 and Nab6 binding across individual transcripts (Fig. 1F and S2B). Each protein mostly targeted the 3’ UTR region, usually at a single site.

Interestingly, the Mrn1 and Nab6 binding sites sometimes coincided (e.g., *HSP150* and *SED1*). There is no reported evidence for direct physical interaction between Mrn1 and Nab6 (Saccharomyces Genome Database), suggesting that the two proteins may compete for binding to some target transcripts.

### Loss of Nab6 confers sensitivity to cell wall damage

Given their clear association with cell wall-related mRNAs, we tested whether disrupting Mrn1 and/or Nab6 function sensitizes yeast to cell wall damage. Two major signaling pathways detect and respond to cell wall damage (Fig. 2A): signals induced by the anti-fungal agents caspofungin, Congo Red (CR), and Calcofluor White (CFW) converge on the CWI MAP-kinase cascade that includes Bck1 and Slt2. Another damage sensor, experimentally triggered by the cell wall-degrading enzyme mixture termed zymolyase elicits a response via CWI and the HOG pathway (Garcia *et al.*, 2015; Sanz *et al.*, 2022).

**Figure 2.**
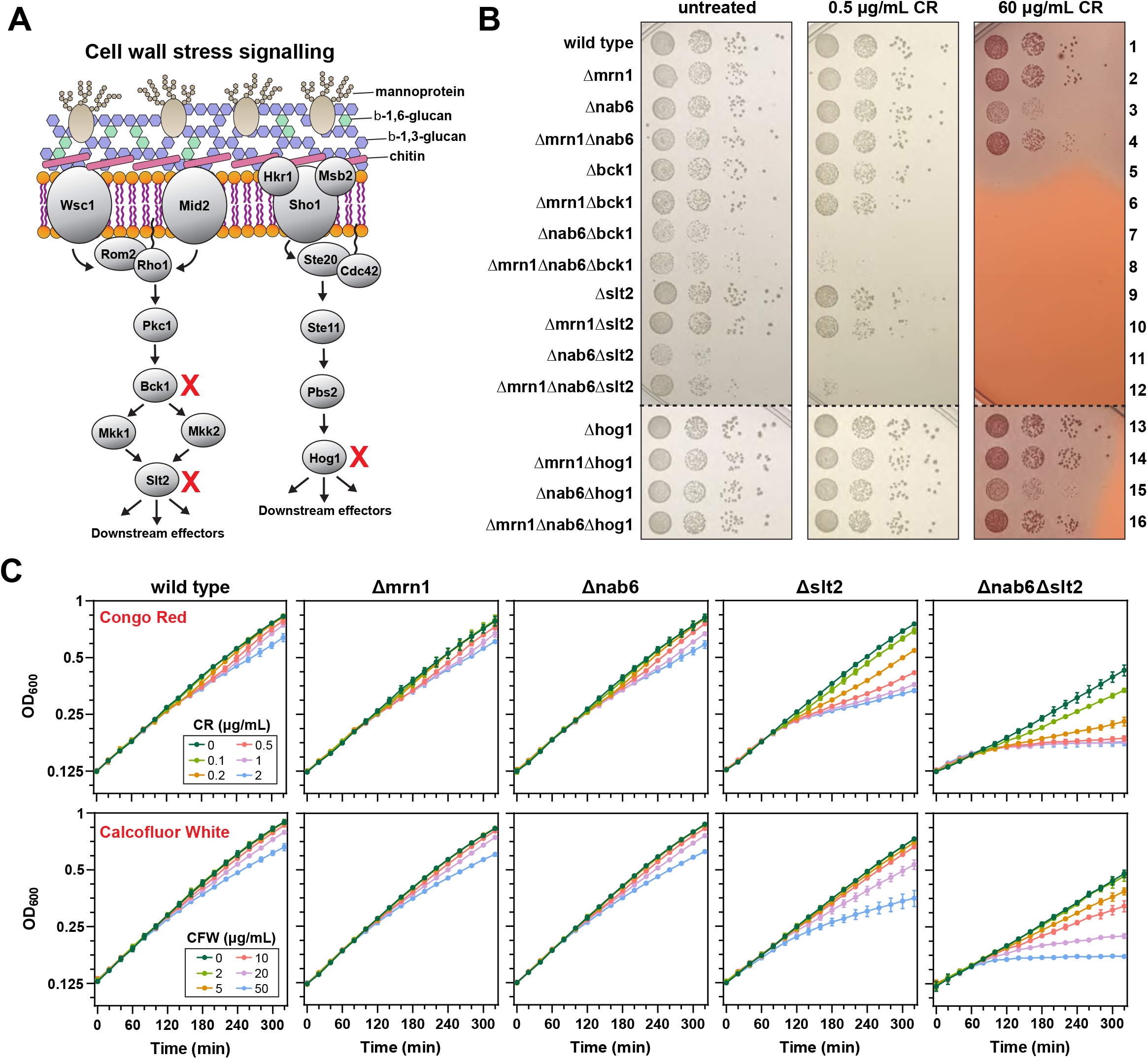
Nab6 acts in parallel to CWI signaling. (A) Schematic outline of the major cell wall stress signaling pathways. The cell wall integrity (CWI) pathway *(left)* responds to Congo Red and Calcofluor White, while the HOG pathway *(right)* responds to zymolyase (Sanz *et al.*, 2022). Genes deleted in (B-C) are marked with a red X. (B) Strains with combinations of gene deletions were tested for their ability to grow on 0.5 μg/mL and 60 μg/mL Congo Red. Cells were grown for two days at 30°C. (C) Growth curves for wild type, *Δmrn1, Δnab6, Δslt2, and Δnab6 Δslt2* strains following treatment with increasing concentrations of Congo Red (*top*) or Calcofluor White *(bottom).* Each point shows the mean (n=2), and error bars indicate standard deviation.

We generated *Δmrn1* and *Δnab6* single mutants and a *Δmrn1 Δnab6* double deletion strain, either alone or in combination with deletions in the CWI *(Δslt2* or *Δbck1)* and HOG *(Δhog1)* pathways (Fig. 2A). Individual deletion strains grew normally, but loss of both Nab6 and CWI pathway components *(Δnab6 Δbck1* and *Δnab6 Δslt2)* gave a clear synthetic growth defect (Fig. 2B). To examine whether Nab6 and/or Mrn1 are involved in cell wall stress response, each strain was challenged with low and high doses of the cell wall-damaging agent CR (Fig. 2B). Low-dose CR (0.5 μg/mL) did not impair growth of the individual knockout strains, but was lethal for the *Δnab6 Δbck1* and *Δnab6 Δslt2* strains. To confirm that the cells were inviable due to compromised cell wall integrity, we repeated the experiment using growth medium supplemented with 1M sorbitol, which provides osmotic support to prevent cell lysis. Notably, sorbitol completely rescued the hypersensitivity of the *Δnab6 Δbck1* and *Δnab6 Δslt2* strains to CR (Fig. S3).

At higher concentrations of CR (60 μg/mL), inactivation of the CWI pathway alone was lethal unless the cells were provided with osmotic support (Fig. 2B and S3, compare rows 1 with 5 and 9). By contrast, the *Δnab6* strain was viable, but substantially growth-impaired (Fig. 2B, compare row 1 and 3). Simultaneous deletion of Nab6 and the CWI pathway *(Δnab6 Δslt2* and *Δnab6 Δbck1)* was lethal with 60 μg/mL CR, and could not be rescued with sorbitol (Fig. 2B and S3A, rows 7 and 11).

Somewhat surprisingly, the individual *Δmrn1* knockout showed no discernable phenotype following CR treatment (Fig. 2B, compare rows 1 and 2). However, *MRN1* showed a clear epistatic relationship with *NAB6.* Deletion of *MRN1* partially rescued the slow-growth phenotype of the *Δnab6* strain at high-dose CR (Fig. 2B, compare rows 3 and 4, and 15 and 16), as well as the lethality observed for the *Δnab6 Δbck1* and *Δnab6 Δslt2* strains at low-dose CR (Fig. 2B, compare rows 7 and 8, and 11 and 12).

Next, we repeated the growth tests in the presence of caspofungin (CSF), an antifungal drug which inhibits the activity of β-1,3-glucan synthase (Szymanski *et al.*, 2022). The individual *Δmrn1* and *Δnab6* strains grew normally, while inactivation of the CWI pathway *(Δbck1* or *Δslt2)* gave a slight growth defect (Fig. S4). However, loss of both Nab6 and the CWI pathway *(Δnab6 Δbck1* and *Δnab6Δslt2)* resulted in a severe synthetic growth defect. Importantly, growth was restored when the medium was supplemented with 1 M sorbitol, confirming that the source of the growth defect was compromised cell integrity.

To analyze growth kinetics in greater detail, we repeated some of the above experiments in liquid medium using various concentrations of CR (Fig. 2C). Wild type cells and individual knockouts were moderately sensitive to CR, while the *Δnab6 Δslt2* strain was inviable at CR concentrations ≥0.5 μg/mL. Notably, the response to CR was not immediate. Wild type cells and individual knockouts were modestly impaired beginning ~100-120 min post-treatment, while the *Δnab6 Δslt2* strain was inhibited by ~80 min. We attribute this to the progressive accumulation of cell wall damage during growth and division. We observed similar growth defects when cells were treated with CFW, an alternative cell wall damaging agent. The individual *Δmrn1* and *Δnab6* strains responded similarly to wild type, while the *Δslt2* strain was moderately inhibited at higher doses (Fig. 2C, *bottom).* By contrast, the *Δnab6 Δslt2* strain was completely inviable at concentrations >50 μg/mL.

Based on these results we propose that Nab6 acts in parallel to the CWI pathway to maintain resistance to the increased cell wall stress generated by CR, CFW, and CSF. The more severe phenotype observed for *Δslt2* compared to *Δnab6* suggests that the CWI pathway mediates the primary response, while Nab6 plays an important supporting role. The related Mrn1 protein has overlapping RNA targets with Nab6 and a partially antagonistic role in responding to cell wall stress caused by CR.

### Nab6 and Slt2 are required for full expression of cell wall mRNAs

Using the growth assays as a guide, we prepared a series of RNAseq libraries from cells exposed to Congo Red (Fig. 3A). Poly(A) selected RNA from wild type, *Δmrn1, Δnab6, Δslt2*, and *Δnab6 Δslt2* cells was harvested under standard conditions (‘time 0’) or following treatment with low-dose CR (2 μg/mL) for 1,2, and 4 hr. In parallel, we treated wild type, *Δmrn1, Δnab6*, and *Δmrn1 Δnab6* cells with high-dose CR (60 μg/mL) and harvested RNA after 2 hr. Overall, replicate libraries were highly reproducible (Fig. S5).

**Figure 3.**
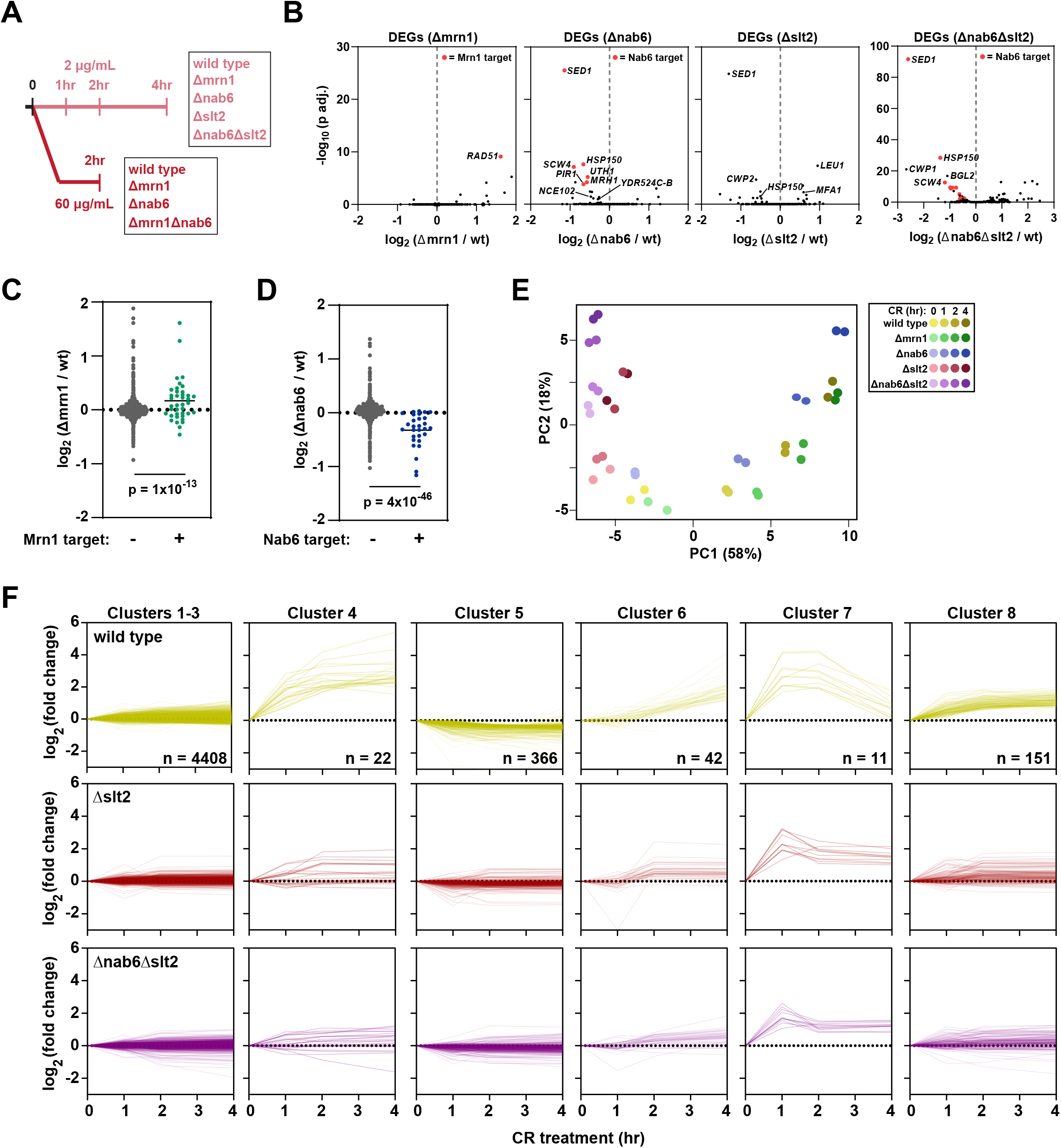
Nab6 positively regulates target mRNAs. (A) Schematic showing the overall design of the transcriptomic experiments. See text for details. (B) Volcano plots showing differentially expressed genes between wild type cells and various knockout strains. Transcripts which are significantly altered (p<0.01) and also a CRAC target of either Mrn1 or Nab6 are colored red. (C) Scatter plots showing the difference in transcript levels between *Δmrn1* and wild type cells. Transcripts are divided into Mrn1 CRAC targets (right) and non-targets (left). The solid black lines show the mean change in transcript levels. The p-value was calculated using a two-sample *t*-test. (D) Same as (C) but for Nab6. (E) PCA for different replicates, timepoints, and strains. (F) Clustering of transcript expression profiles for wild type cells following CR treatment. Expression profiles for the same genes are shown below for the *Δslt2* and *Δslt2 Δnab6* strains.

In untreated cells (Fig. 3B) only four transcripts were differentially expressed between *Δmrn1* and wild type cells. The most significantly upregulated transcript, *RAD51*, was also a strong Mrn1 binding target. (highlighted in red in Fig. 3B). As a group, Mrn1 CRAC targets were mildly upregulated compared to non-targets, suggesting reduced stability from Mrn1 binding (Fig. 3C). In the *Δnab6* strain, 9 transcripts were significantly downregulated, and 6 of these were also strong Nab6 CRAC targets (Fig. 3B). The most significantly altered transcript was *SED1*, encoding a major stress-induced structural component of the cell wall (Shimoi *et al*, 1998). Overall, Nab6 targets were substantially reduced in abundance compared to non-target transcripts in *Δnab6* (Fig. 3D), indicating that Nab6 binding results in mRNA stabilization.

The *Δslt2* strain had relatively few differentially expressed genes compared to the wild type strain (Fig. 3B). As with *Δnab6*, the most significantly altered transcript was *SED1*, but several additional cell wall mRNAs were also downregulated, including *BGL2*, encoding Endo-beta-1,3-glucanase, a key enzyme in cell wall synthesis and remodeling. The combined *Δnab6 Δslt2* strain showed a further decrease in cell wall mRNAs compared to either single deletion (Fig. 3B), suggesting that Nab6 and Slt2 act independently to maintain normal levels of cell wall mRNAs.

We next determined the effects of Congo Red. To our surprise, the different CR treatments (2 and 60 μg/mL) induced nearly identical changes in the transcriptome (Fig. S6). We therefore focused on the 2 μg/mL dose for subsequent analysis. Initially, we used principal component analysis (PCA) to compare the global mRNA expression patterns for each strain throughout the time course (Fig. 3E). The wild type and *Δmrn1* strains showed similar profiles; *Δnab6* was more distinct but followed the same general trend. By contrast, the *Δslt2* strain showed a substantially altered response to CR, and this was even more marked in the *Δnab6 Δslt2* strain. To further investigate this observation, we divided the mRNAs into eight clusters based on their response to CR in wild type cells (Fig. 3F, Table S3). Transcripts from clusters 1-3 (n=4408) were largely unchanged throughout the time course and were grouped together for subsequent analyses. The remaining mRNAs (n=592) fell into one of five clusters based on their response profile. Cluster 5 mRNAs (n=366) were modestly decreased, while mRNAs in clusters 4, 6, and 8 (n=215) were all upregulated, but with varying kinetics. Cluster 7 transcripts (n =11) were strongly upregulated between 0 and 1 hr post treatment, plateaued between 1 and 2 hr, and decreased thereafter. *SLT2* deletion markedly reduced, but did not completely abolish, the changes in transcript levels observed in clusters 4 through 8 (Fig. 3F, *middle* and *lower* panels). Additional loss of Nab6 further muted the response to CR. This was particularly apparent for the strong Nab6 targets *HSP150* and *SED1* (Figs. S7A-B).

The *Δnab6 Δslt2* strain cannot grow in the presence of low-dose CR, even though the individual deletion strains are viable (Fig. 2). To understand the molecular basis for this phenotype, we compared the transcriptomic profile of the Δ*nab6* Δ*slt2* double mutant with the Δ*slt2* single mutant (Fig. 4). Several transcripts were downregulated in Δ*nab6* Δ*slt2* cells compared to Δ*slt2*, most of which were also direct binding targets for Nab6 (highlighted in red). Overall, these results are consistent with previous reports that the CWI pathway plays a major role in transcriptional reprogramming during cell wall stress (Boorsma *et al*, 2004; Garcia *et al.*, 2004) but support a parallel function for Nab6.

**Figure 4.**
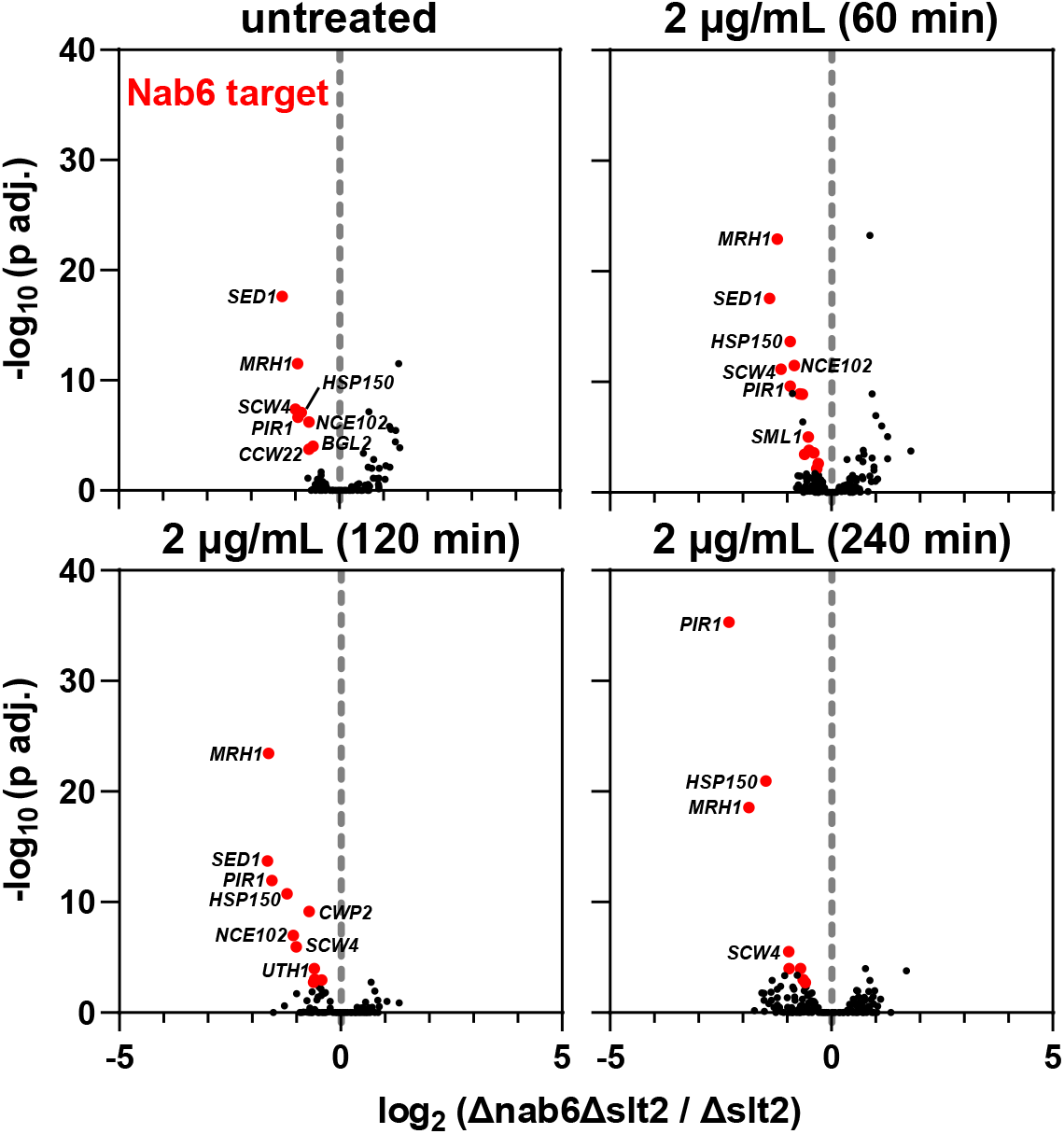
Loss of SLT2 exacerbates the expression defects associated with *Λnab6*. Volcano plots showing differentially expressed genes between *Δnab6 Δslt2* knockout strains and the *Δslt2* single mutant. Transcripts which are significantly altered (p<0.01) and also a CRAC target of Nab6 are colored red.

### Loss of Mrn1 partially rescues molecular defects associated with the *Δnab6* strain

The growth tests revealed that Mrn1 is partially epistatic to Nab6 upon treatment with CR (Fig. 2). To identify a molecular basis for this phenotype, we compared the *Δmrn1 Δnab6* double mutant with the *Δnab6* single mutant, in the absence or presence of high dose CR. Only a small number of transcripts were differentially expressed between the two strains (Fig. 5A-B). However, two of these mRNAs *(SED1* and *BGL2)* encode important cell wall components and are bound by Nab6 and Mrn1 (Table S1). In unstressed cells, both transcripts were downregulated in the absence of Nab6, but mRNA levels were restored in the double mutant. CR treatment increased the abundance of *SED1* and *BGL2* (note differences in scales), but to a lesser extent in the *Δnab6* strain (Fig. 5C). Notably, additional deletion of Mrn1 once again rescued expression of *SED1* and *BGL2.* These findings support the model that Nab6 and Mrn1 have opposing roles in regulating the abundance of mRNAs encoding major cell wall proteins.

**Figure 5.**
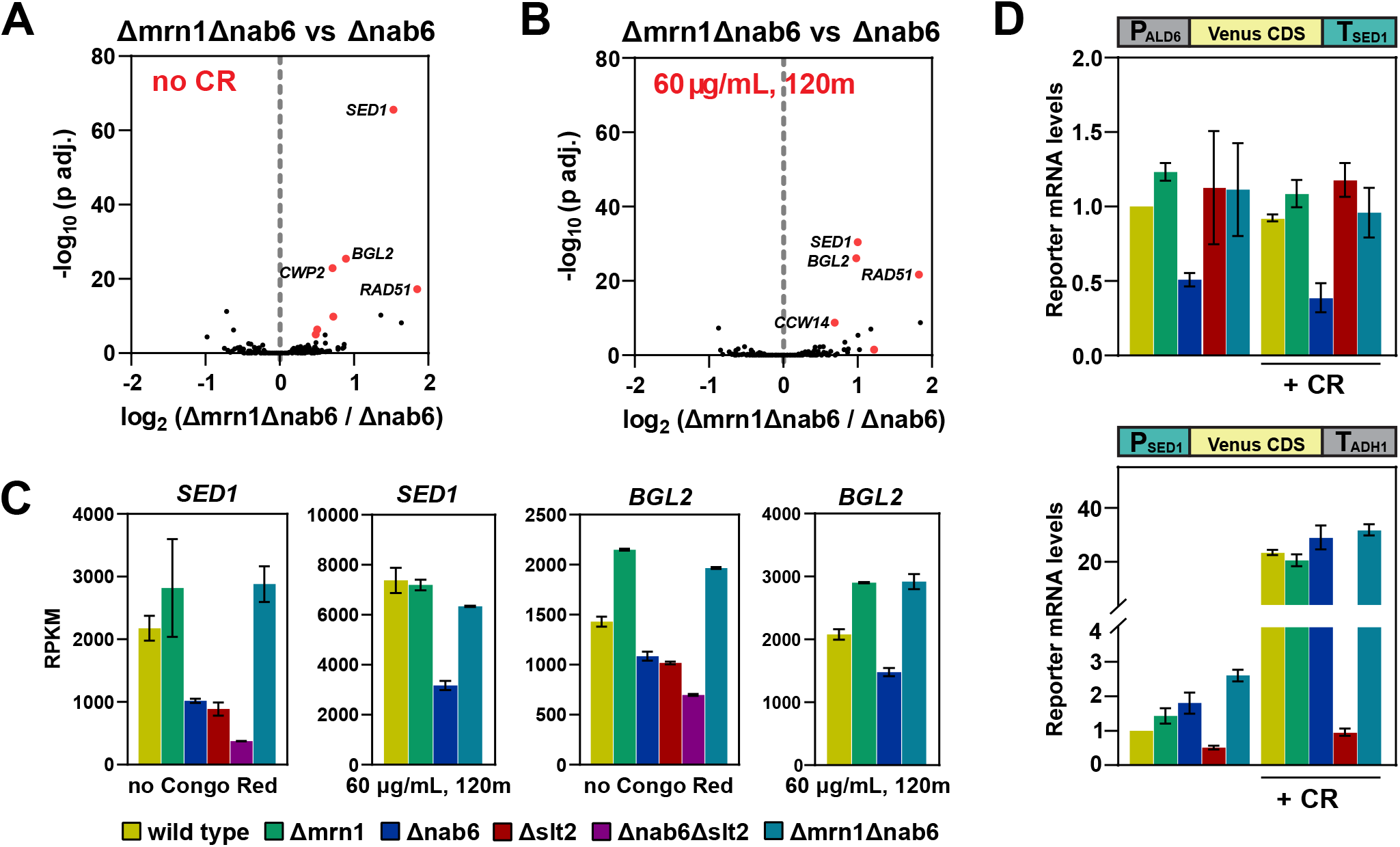
Loss of Mrn1 partially suppresses the phenotype from loss of Nab6. (A-B) Volcano plots showing differentially expressed genes between *Δmrn1 Δnab6* knockout strains and the *Δnab6* single mutant. Transcripts which are significantly altered (p<0.01) and also a CRAC target of either Mrn1 or Nab6 are colored red. (C) Bar graphs showing the number of reads (fragments per kilobase per million mapped reads) in each library which map to the *SED1* or *BGL2* transcripts in the absence or presence of CR. Error bars show standard deviation of the mean (n=2). (D) Bar graphs showing the mRNA levels for the T_SED1_ (*top*) and P_SED1_ *(bottom)* reporter constructs. Error bars show standard deviation of the mean (n=2). Bar colors are the same as in (C).

The predominant binding of Nab6 and Mrn1 to mRNA 3’ UTRs, including that of *SED1*, suggested that this region would be required for regulation by these proteins. To test this, we generated a reporter construct in which the coding sequence (CDS) for the fluorescent protein Venus was flanked by the promoter and 5’ UTR from *ALD6* (P_ALD6_) and the 3’ UTR and terminator sequence from *SED1* (T_SED1_). P_ALD6_ was chosen because the *ALD6* mRNA is expressed at similar levels to *SED1* and is not altered in response to CR treatment (Table S2). The resulting construct was chromosomally integrated and mRNA expression was assessed by RT-qPCR. The T_SED1_ reporter mRNA showed a marked reduction in expression in the absence of Nab6 (*Δnab6*), which could be rescued by additionally deleting *MRN1* (Δ*mrn1* Δ*nab6*) (Fig. 5D, *top*). These data support the model that Nab6 and Mrn1 have opposing effects on *SED1* mRNA abundance, acting posttranscriptionally via the 3’ UTR.

For comparison, we generated a second reporter construct consisting of the promoter and 5’ UTR from *SED1* (P_SED1_) fused to the Venus CDS and the 3’ UTR and terminator sequence from *ADH1* (T_ADH1_), which is not stress responsive. This construct is expected to be under transcriptional control of CWI, while avoiding Nab6/Mrn1 post-transcriptional regulation. CR treatment induced a robust increase in expression that was abrogated by deletion of *SLT2* (Fig. 5D, *bottom*), confirming that the authentic transcriptional regulation has been recapitulated. By contrast, induction was not affected by loss of Nab6 or Mrn1, consistent with these proteins acting posttranscriptionally.

## DISCUSSION

An estimated ~180 genes are directly involved in cell wall biosynthesis, remodeling, or structure (Orlean, 2012). Many encode proteins that are directly incorporated into the cell wall, but most regulate upstream processing events, such as precursor synthesis, O-mannosylation, N-glycosylation, or GPI attachment. Our CRAC experiments revealed that Mrn1 and Nab6 predominantly target mRNAs from the former group; examples include both structural components of the wall (e.g. *HSP150* and *SED1)* and remodeling enzymes (e.g. *SCW4, BGL2, DSE2*, and *CTS1*). Typically, these transcripts were bound by Mrn1/Nab6 at only one or two sites within their 3’ UTRs, indicating highly specific binding.

Both Mrn1 and Nab6 also targeted additional mRNAs not directly implicated in cell wall biosynthesis. These included several mRNAs encoding plasma membrane proteins (Pmp1, Pma1, Mrh1, and Nce102), and the mating pheromone Mfa1. These proteins are matured through the same general secretory pathway as cell wall proteins (Coïc *et al*, 2005; Grossmann *et al*, 2008; Luo & Chang, 2000; Michaelis & Barrowman, 2012; Wu *et al*, 2000); reviewed in (Feyder *et al*, 2015; Free, 2013), possibly explaining why seemingly unrelated mRNAs might be regulated by the same RNA-binding proteins.

One of the most enriched Mrn1-specific targets was the *MRN1* transcript itself (Table S1), suggesting possible autoregulation. Notably, Mrn1 binding to *MRN1* was situated entirely within the 5’ UTR, a pattern contrary to all other major targets (Fig. S2B). We speculate that binding to the *MRN1* 5’ UTR might reduce stability and/or inhibit translation, leading to autoregulation.

Most Nab6 targets showed decreased expression in the *Δnab6* strain, and even more so in *Δnab6 Δslt2*, suggesting that Nab6 stabilizes its target mRNAs. The expression changes were relatively modest, but phenotypically significant. In the absence of the CWI pathway, Nab6 was completely essential for cells to survive even low levels of CR or CFW. Even in the absence of specific cell wall stress, the *Δnab6 Δslt2* strain was slow-growing, suggesting Nab6 also plays a role in normal cell wall integrity and/or the remodeling needed for cell division.

Only four transcripts were differentially expressed in the absence of Mrn1, but one of these *(RAD51)* was also a direct mRNA binding target for Mrn1. These results are broadly consistent with a recent study which analyzed the transcriptome of *Δmrn1* cells (Reynaud *et al*, 2021). The same study also reported that Mrn1 recruits the cellular RNA degradation machinery, consistent with our observation that at least some Mrn1 targets are upregulated in the *Δmrn1* strain. However, we cannot exclude the possibility that Mrn1 regulates additional aspects of gene expression, such as mRNA localization or translation.

An interesting phenotype emerged when we compared the *Δnab6* strain with *Δmrn1 Δnab6* double mutants. *SED1* and *BGL2* were downregulated in the absence of Nab6, but mRNA levels were rescued when Mrn1 was also deleted. Both transcripts were bound by Nab6 and Mrn1 at overlapping sites, supporting a model in which the two proteins compete for binding. In the absence of Nab6, Mrn1 may show enhanced binding, destabilizing the mRNA; if both proteins are absent, then mRNA expression is restored. We were initially surprised that Mrn1 and Nab6 have antagonistic functions, at least on some cell wall-related substrates. However, the cell wall must constantly be remodeled to allow for expansion during growth, while retaining integrity against turgor pressure, even in the absence of specific damage. So, an accurate balance between activities that weaken and strengthen the cell wall must always be maintained. We speculate that an interplay between Nab6 and Mrn1 activities aids this process.

Previous studies of cell wall stress have largely focused on the CWI signaling pathway. In response to cell wall damage, the CWI pathway broadly activates the transcription of cell wall-related mRNAs. Phenotypic changes induced by transcriptional regulation are generally strong, but also relatively slow; induced mRNAs must be transcribed, processed, and exported prior to translation. Posttranscriptional regulation bypasses some or all of these steps, potentially eliciting a much faster response. Posttranscriptional control also allows fine-tuning; mRNAs can be localized and translated only when and where they are needed, e.g. at sites of cell wall growth or repair. Ssd1 appears to function precisely in this way, repressing translation of mRNAs until they are delivered to the bud neck (Jansen *et al.*, 2009). Nab6 may have an analogous role, stabilizing and/or activating translation of mRNAs destined for sites of growth or repair.

Recent work from the Chanfreau lab highlighted the importance of posttranscriptional regulation for cell wall homeostasis (Wang *et al*, 2020). The *Δslt2* strain is severely impaired in response to heat shock, a condition which also induces cell wall stress (de Nobel *et al*, 2000; Wang *et al.*, 2020). However, additional deletion of various individual mRNA decay factors (e.g. the exosome, Xrn1, or Upf1), was sufficient to restore growth (Wang *et al.*, 2020). This observation suggests RNA synthesis and degradation are in balance. To survive cell wall stress, cells must either transcribe new cell wall mRNAs or stabilize the existing pool. By protecting its target mRNAs, Nab6 may partially compensate for the absence of transcriptional regulation in *Δslt2* cells. Only in the absence of both Slt2 and Nab6 do cells become especially vulnerable to cell wall damage.

In summary, our results implicate Nab6 and Mrn1 as key regulators of cell wall homeostasis. We showed that Nab6 and Mrn1 act in parallel to conventional CWI signaling to maintain adequate cell wall mRNA expression in response to cell wall damage. However, important questions remain unresolved. What is the mechanism of target mRNA stabilization by Nab6? Is this its only function, or does Nab6 also regulate mRNA localization and/or translation? More broadly, why do so many RBPs specifically target cell wall mRNAs (Bayne *et al.*, 2022; Hogan *et al.*, 2008; Tuck & Tollervey, 2013)? Do they act independently or in concert? The cell wall is an important target for several clinically-approved antifungal agents, but their use is limited by fungal resistance (Chandrasekar, 2007; Eschenauer *et al*, 2007; Wagener & Loiko, 2017). Identifying the mechanisms by which fungi respond to cell wall stress will ultimately result in improved treatment options.

## DATA AVAILABILITY

The sequencing data generated in this study can be accessed on the Gene Expression Omnibus (GEO) with accession number GSE210558.

## ACKNOWLEDGEMENTS

We thank Richard Clark, Audrey Coutts, and Angie Fawkes from the Edinburgh Wellcome Clinical Research Facility for sequencing services as well as Shaun Webb from the WCB bioinformatics core facility for maintaining the servers we used for processing sequencing data. This work was funded by Wellcome Principal Research Fellowships to DT (109916 and 222516), also supporting SB and VS, and core funding for the Wellcome Centre for Cell Biology (203149).

## AUTHOR CONTRIBUTIONS

Conceptualization: SB, VS, and DT. Experiments: SB, VS. Data analysis: SB. Writing – Original Draft: SB. Writing – Review and Editing: SB, DT, VS. Funding Acquisition: DT.

## DECLARATION OF INTERESTS

The authors declare no competing interests.

## STAR METHODS

### Cell culture and medium

All yeast strains were cultured at 30°C in synthetic dropout (SD) -trp -met supplemented with 20 μg/mL methionine and 2% glucose. CRAC samples were crosslinked at 0.4 OD_600_. Growth curves and RNAseq experiments were initiated at 0.125 OD_600_. For the 10-fold serial dilution assays, cells were grown to stationary phase overnight, diluted to 0.5 OD_600_, and spotted on -trp plates supplemented with varying concentrations of Congo Red or caspofungin.

### Gene tagging and deletion

For CRAC experiments, the chromosomal copies of MRN1 and NAB6 were C-terminally tagged with HF (His-Flag), consisting of eight consecutive histidine residues, a four alanine spacer, and a single Flag motif (HHHHHHHHAAAADYKDDDDK). The two proteins were tagged using CRISPR-Cas9 and the pML104 vector (Laughery *et al*, 2015) as described below.

The pML104 vector included a Cas9 expression construct, a URA3 selectable marker, and a guide RNA (gRNA) cloning site. Approximately 10 μg of plasmid was digested overnight with SwaI (NEB Cat#R0604S), and then for 2 hr at 50°C with BclI-HF (NEB Cat#R3160S). The digested vector was purified by gel extraction, and aliquoted for later use.

Guide RNA oligos were designed as previously reported (Laughery *et al.*, 2015). Each oligo pair was annealed in a reaction consisting of 1 μM forward oligo, 1 μM reverse oligo, and 1X T4 DNA ligase buffer (NEB Cat#B020S) in a 100 μL reaction volume. The hybridization reaction was initially incubated at 95°C for 6 min, and gradually decreased to 25°C at the rate of 1.33°C/min. Hybridized substrates were then ligated into the digested vector at 25°C for 4 hr. The ligation reaction consisted of 265 ng pre-cut pML104 vector, 0.8 nmol insert, 1X T4 DNA ligase buffer, and 800U of T4 DNA ligase (NEB M0202L) in a 40 μL reaction volume. The ligation mix was transformed into homemade DH5a E. coli, and plated overnight on LB-Amp. Plasmid DNA was isolated and sequenced to ensure correct insertion of the guide sequence.

To insert the HF tag, we designed fragments consisting of the HF DNA sequence flanked by 50 bp homology arms to serve as repair templates. Synonymous mutations were typically incorporated into each construct in order to disrupt the PAM site and prevent further cleavage by Cas9. Each repair template was made by annealing two single-stranded oligonucleotides sharing 20 bp of complementarity at their 3’ ends. Each set of oligos was annealed in a reaction consisting of 10 μM forward oligo, 10 μM reverse oligo, and 1X NEB buffer 2.1 in a 43 μL reaction volume. The hybridization reaction was initially incubated at 95°C for 6 min, and gradually decreased to 25°C at the rate of 1.33°C/min. Subsequently, the annealed oligos were incubated in the same buffer supplemented with 250 μM dNTPs (Takara Cat#RR002M) and 5U Klenow exo- (NEB Cat#M0212L) in a 50 μL reaction at 37°C for 1 h to fill in the single-stranded regions. To introduce the HF tag, BY4741 yeast were transformed using the standard PEG/LiOAc protocol with 500 ng of gRNA plasmid and 10 pmol of the corresponding repair template. Transformants were plated onto -ura medium. After three days, several clones from each transformation were plated again on selective medium, and allowed to grow for an additional 2-3 days. Single colonies were selected and plated on YPD for 2 days. Finally, individual colonies were grown overnight in liquid YPD and frozen. The clones were verified by PCR using flanking primers and confirmed by sequencing.

With the exception of HOG1, all gene deletions were also made using CRISPR/Cas9. Guide RNA plasmids and repair templates were generated essentially as described above. The *Δhog1* strain was generated with conventional homologous replacement using a HIS3 selectable marker derived from pFA6a-HIS3MX6 (Addgene: 41596).

### CRAC

The CRAC protocol was performed as previously described (Bresson *et al.*, 2020). For each CRAC experiment, 700 mL of cells were cultured in SD -trp medium. At OD_600_ 0.4 the cells were UV-irradiated at 254 nm with a dose of 100 mJ/cm^2^ (4-6 s) using the Vari-X-Link crosslinker. Following crosslinking, cells were collected by filtration and resuspended in 50 mL of ice-cold PBS, and centrifuged at 4600g for 2 min. The cell pellets were frozen and stored at −80°C. Subsequently, cell pellets were resuspended in 500 μL TN150 (50 mM Tris-HCl pH 7.5, 150 mM NaCl, 0.1% NP-40, 5 mM β-mercaptoethanol and a protease-inhibitor cocktail (1 tablet / 50 mL) (Roche Cat#11873580001). The resuspended cells were added to 1.25 mL zirconia beads in a 50 mL conical and lysed using 5 one-minute pulses, with cooling on ice in between. The resulting lysate was further diluted with 1.5 mL TN150, briefly vortexed, and centrifuged at 4600g for 5 min. The supernatant was transferred to a 1.5 mL Eppendorf tube and centrifuged at 16000g for 20 min. In parallel, 100 μL of magnetic anti-Flag bead slurry (Sigma-Aldrich Cat#M8823) was washed twice with TN150. The cleared lysate was incubated with the anti-Flag beads at 4°C with nutating. After 2 hr, the supernatant was removed, and the beads were washed four times with TN150 (5 min nutating at 4°C for each wash). To elute the protein of interest, the beads were incubated with 20 μg of Flag peptide (Sigma-Aldrich Cat#F3290) in a 200 μL volume at 37°C for 15 min with shaking. The eluate was transferred to a fresh tube containing 350 μL TN150 and treated with RNace-IT (Agilent Cat#400720) (0.1U, 5 min, 37°C) to fragment RNA. The RNase reaction was halted by transferring the eluate to a tube containing 400 μg guanidine hydrochloride. The solution was adjusted for nickel affinity purification with the addition of 27 μL 5 M NaCl and 3 μL 2.5 M imidazole, and added to 50 μL of washed nickel beads (QIAGEN Cat#30410). After overnight nutation at 4°C, the beads were transferred to a spin column (Thermo Scientific Cat #69725), washed 3 times with 400 μL WBI (6 M guanidine hydrochloride, 50 mM Tris-HCl pH 7.5, 300 mM NaCl, 0.1% NP-40, 10 mM imidazole, and 5 mM β-mercaptoethanol), and then 3 times with 600 μL PNK buffer (50mM Tris-HCl pH 7.5, 10 mM MgCl_2_, 0.5% NP-40, and 5 mM β-mercaptoethanol). Four subsequent reactions (80 μL each) were performed on-column, and subsequently washed once with WBI and three times with PNK buffer:

1. Phosphatase treatment (1x PNK buffer, 8U TSAP (Promega, Cat#M9910), 80U RNaseIN (Promega Cat#N2511); 37°C for 30 min).
2. 3’ linker ligation (1x PNK buffer, 20U T4 RNA ligase I (NEB Cat#M0204L), 20U T4 RNA ligase II truncated K227Q (NEB Cat#M0351L), 80U RNaseIN, 1 μM Preadenylated 3’ miRcat-33 linker (IDT); 25°C for 6 hr).
3. 5’ end phosphorylation and radiolabeling (1x PNK buffer, 40U T4 PNK (NEB Cat#M0201L), 40 μCi ^32^P-γATP; 37°C for 60 min, with addition of 100 nmol ATP after 40 min).
4. 5’ linker ligation (1x PNK buffer, 40U T4 RNA ligase I, 80U RNaseIN, 5’ linker, 1 mM ATP; 16°C overnight).

After the final ligation reaction, the beads were washed twice with WBII (50 mM Tris-HCl pH 7.5, 50 mM NaCl, 0.1% NP-40, 10 mM imidazole, and 5 mM β-mercaptoethanol). Protein:RNA complexes were eluted twice (10 min each) in 40 μL elution buffer (same as WBII but with 300 mM imidazole). At this point, different replicates for the same protein were combined. The merged eluates were precipitated with 5X volume acetone at −20°C for at least two hours. Protein:RNA complexes were pelleted (16000g, 20 min, 4°C) and resuspended in 20 μL 1X NuPAGE sample loading buffer (Invitrogen Cat#NP0007) supplemented with 8% β-mercaptoethanol. The sample was denatured by incubation at 65°C for 10 min and run on a 4-12% Bis-Tris NuPAGE gel (Invitrogen Cat#NP0321BOX) at 150V in 1X NuPAGE MOPS buffer (Invitrogen Cat#NP001-02). The protein:RNA complexes were transferred to Hybond-N+ nitrocellulose membranes (GE Healthcare Cat#RPN303B) with NuPAGE transfer buffer (Invitrogen Cat#NP0006-1) for 1.5 hr at 100V, and detected using autoradiography. The appropriate region was excised from the membrane and treated with 0.25 mg/mL Proteinase K (Roche Cat#03115836001) for 2 hr at 55°C in a 500 μL reaction consisting of 50 mM Tris-HCl pH 7.5, 50 mM NaCl, 0.1% NP-40, 10 mM imidazole, 1% SDS, 5 mM EDTA, and 5 mM β-mercaptoethanol. The RNA component was isolated using phenol:chloroform extraction followed by ethanol precipitation. Subsequently, the RNA was reverse transcribed using Superscript III (Invitrogen Cat#18080-044) and the miRCat-33 RT oligo (IDT) for 1 hr at 50°C in a 20 μL reaction. The resulting cDNA was amplified by PCR in five separate reactions using La Taq (Takara, Cat#RR002M) (2 μL template in each reaction, 21 cycles). The PCR reactions were combined, precipitated in ethanol, and resolved on a 3% Metaphore agarose gel (Lonza Cat#50180). A region corresponding to 140-200 bp was excised from the gel and extracted (QIAGEN Cat#28606). Libraries were sequenced by the Wellcome Trust Clinical Research Facility (Edinburgh, UK) on Next-seq with single-end, 75-nt output.

### RNAseq

Approximately 6 ODs of cells were harvested by centrifugation (4600 rpm; 1min) and frozen at −80°C. Subsequently, the cells were lysed using zirconia beads (Thistle Scientific Cat#ZrOB05) and RNA was extracted with phenol:chloroform followed by ethanol precipitation. Glycoblue (Fisher Scientific Cat#AM9516) was used as a coprecipitant and to visualize the RNA pellet after centrifugation. Libraries for RNAseq were prepared by the Wellcome Trust Clinical Research Facility at Western General Hospital (Edinburgh, UK) using the poly(A) mRNA magnetic isolation kit (NEB Cat#E7490) and the NEBNEXT Ultra II Directional RNA Library Prep kit (NEB Cat#7760), and then sequenced using Next-Seq with paired-end, 100nt output.

### Reporter constructs

The reporter constructs were designed using the modular cloning system devised by the Dueber lab (Lee *et al*, 2015). In this system, each reporter cassette is comprised of nine different ‘parts’, such as promoters, coding sequence, terminators, etc. A collection of standard parts was obtained from Addgene (Kit# 1000000061), and parts specific to this study were generated using standard PCR and cloning techniques (Lee *et al.*, 2015).

Individual parts were combined into reporter cassettes in a reaction consisting of 50 ng of each part plasmid, 1 μL T4 DNA ligase buffer, 0.5 μL T4 DNA ligase (NEB Cat#M020S), 0.5 μL BsaI-HFv2 (NEB Cat#R3733S), and water to bring the final volume to 10 μL. Reaction mixtures were incubated in a thermocycler for 25 cycles of digestion and ligation (42°C for 2 min, 16°C for 5 min) followed by a final digestion step (60°C for 10 min) and heat inactivation (80°C for 10 min). The assembled plasmids were confirmed via diagnostic restriction digest. Subsequently, the plasmids were linearized by digestion with NotI-HF (NEB Cat#R3189S), and transformed into the desired strains. Clones were selected on YPD supplemented with 200 μg/mL hygromycin (Millipore Cat#400052-50ML) and verified by PCR. Reporter experiments were performed in both technical and biological duplicate. The two technical replicates were averaged together to make one biological replicate.

### qPCR

Approximately 2 ODs of cells were harvested by centrifugation (4600 rpm; 1min), frozen at −80°C, and lysed using zirconia beads. RNA was extracted using phenol:chloroform, and residual DNA was degraded by treatment with Turbo DNase (ThermoFisher Cat#AM2238) in a 50 μL reaction consisting of 10-25 μg of RNA, 1 U DNase, and 20 U of RNasIN in a 1X solution of the manufacturer-supplied buffer at 37°C for 30 min. Subsequently, the DNase was removed by phenol-chloroform extraction followed by ethanol precipitation. Reverse transcription (RT) and qPCR were performed using the Luna Universal one-step RT-qPCR kit (NEB Cat#E3005S) in a 5 μL reaction consisting of 300 pmol primers, 1X reaction mix, 1X enzyme mix, and 8.75 ng of RNA. Reaction mixtures were incubated in a thermocycler with an initial RT reaction (55°C for 10 min) followed by 40 cycles of amplification (95°C for 5s and 60°C for 10s). Each sample was measured in technical quadruplicate and the results were averaged together. Relative expression was calculated using ΔΔC_t_ and by normalizing to the levels of *RPL6B* mRNA in each sample.

### Protein domain analysis

Protein domains were predicted using SMART (http://smart.embl-heidelberg.de/). The RRM structures were aligned using the `super` command in Pymol. Sequence alignments were performed using Clustal Omega (https://www.ebi.ac.uk/Tools/msa/clustalo/).

### CRAC analysis

Bioinformatic analysis of CRAC datasets was performed as previously described (Bresson *et al.*, 2020). Multiplexed CRAC datasets were separated using `pyBarcodeFilter` from the pyCRAC package (Webb *et al*, 2014). Subsequently, Flexbar v3.4.0 (Dodt *et al*, 2012) was used to remove sequencing adapters, trim low-quality baes from the 3’ end, and remove low-quality reads (parameters -ao 4 -u 2 -q TAIL -m 14 -at RIGHT with adapter sequence TGGAATTCTCGGGTGCCAAGGC. In addition to the barcode, each read contained three random nucleotides at the 5’ end to allow PCR duplicates to be collapsed using `pyFastqDupiicateRemover` (Webb *et al.*, 2014). Reads were filtered to exclude low-entropy sequences using `bbduk` (https://sourceforge.net/projects/bbmap/) (parameters: entropy = 0.65 entropywindow = 10 entropyk = 5). Subsequently, the reads were mapped to a modified version of the *S. cerevisiae* EF4.74 genome (Ensembl) in which the introns had been bioinformatically removed (Bresson *et al.*, 2020). Sequencing reads were aligned using `Novoaiign` v2.07.00, with reads mapping to multiple locations randomly assigned (-r Random). Reads which aligned to the same coordinates and had identical 5’ random barcodes were collapsed into a single read.

Genome coverage maps were generated using `genomecov` from `bedtools` v2.27.0 and visualized using IGV (https://software.broadinstitute.org/software/igv/). The number of reads mapping to different mRNAs was tabulated using `pyReadCounters` and a custom genome annotation file. Gene ontology mapping was generated using the Proteomaps website (Liebermeister *et al*, 2014).

### RNAseq analysis

Paired-end reads were aligned to the intronless *S. cerevisiae* genome using `STAR` and tabulated using `featureCounts`. Genome coverage files were `generated` using genomecov from bedtools and scaled by fragments per million. For all analyses, we used the top 5000 transcripts, defined by their average expression across all replicates and conditions. Volcano plots and PCA were performed using R with `DESeq2` and `ggplot2`. The clustering analysis was performed on log_2_ transformed data using Morpheus (https://software.broadinstitute.org/morpheus) (settings: Euclidean distance, 8 clusters).

---

**Figure S1.**
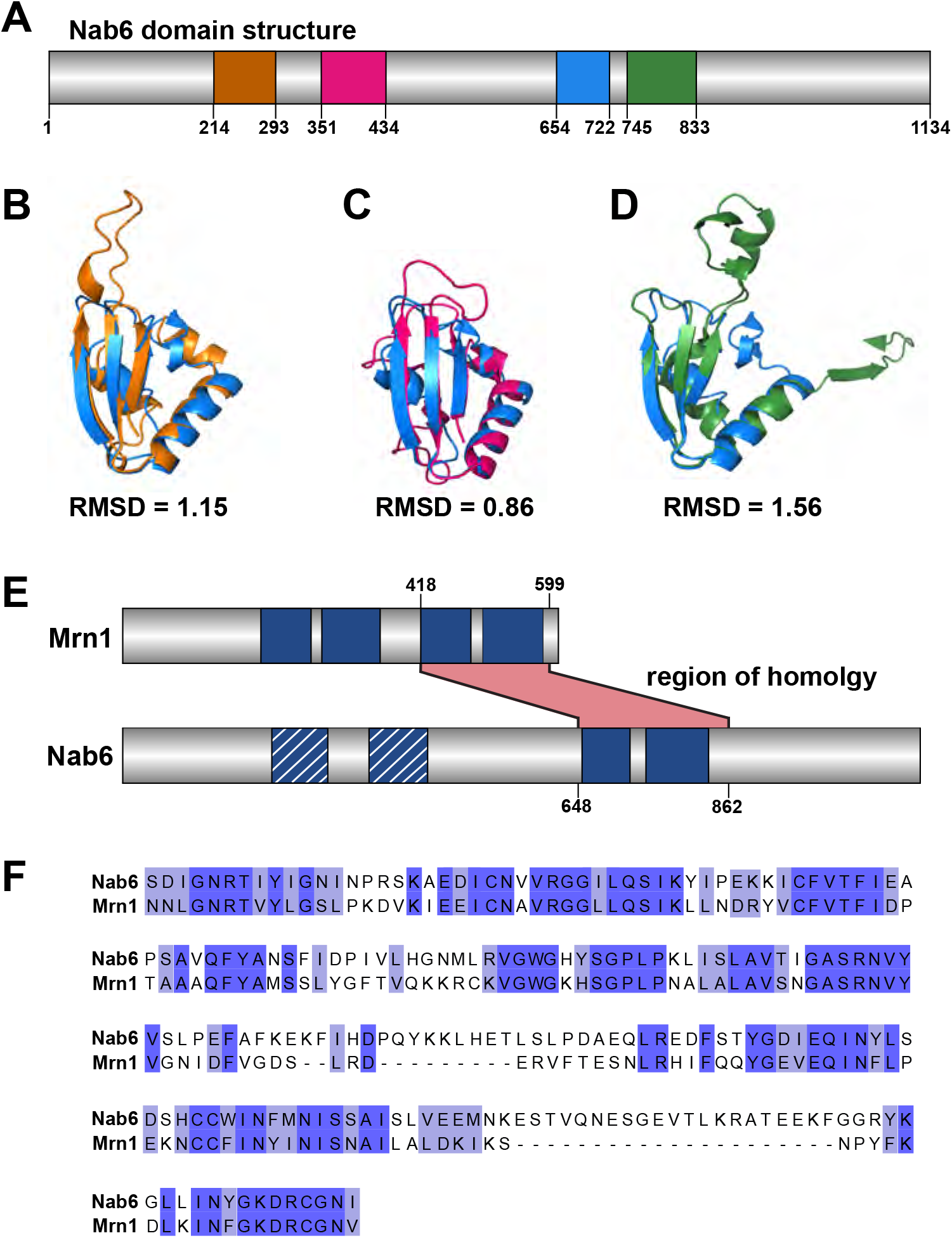
Related to Figure 1. (A) Domain architecture of Nab6 with each RRM shown in a different color. RRM1: 214-293 (orange). RRM2: 351-434 (pink). RRM3: 654-722 (blue). RRM4: 745-833 (green). (B-D) Structural alignments of RRM3 with RRM1 (B), RRM2 (C), and RRM4 (D). The colors are the same as in (A). (E) Domain architecture of Mrn1 and Nab6. Two putative RRMs identified visually using Alphafold are shown in blue with white stripes. The region of homology between Mrn1 and Nab6 is indicated in pink. (F) Sequence alignment of the region of homology between Mrn1 and Nab6. Identical residues are shown in dark blue, and similar residues are shown in light blue.

**Figure S2.**
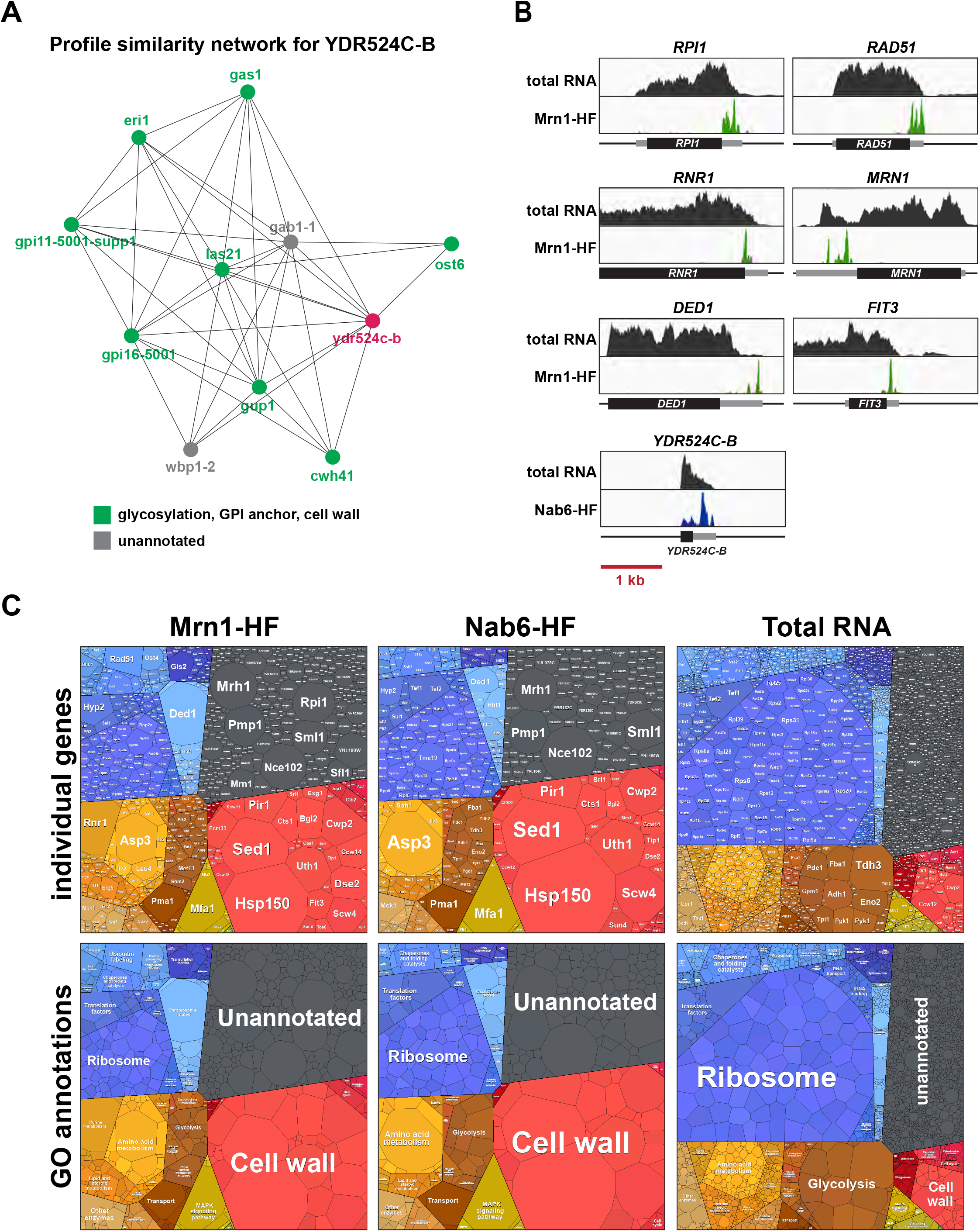
Related to Figure 1. (A) Genetic profile similarity map for YDR524C-B showing genes with related sets of genetic interactions. Genes with a known role in cell wall biogenesis are marked in green. (B) Binding of Mrn1 and Nab6 across selected target transcripts preferentially targeted by one protein or the other. Each track is normalized to total library size using reads per million. RNAseq reads are included as a control. Each box represents a 3 kb window; a scale bar is shown at the bottom. The open reading frames (ORFs) are shown as black boxes, with UTRs as flanking gray boxes. Each transcript is oriented 5’ to 3’. (C) Area plots showing the proportion of each library mapping to individual genes. The upper panels show individual genes, and the lower panels show GO terms. Note that because of the way the data is represented, multifunctional genes can only be assigned to a single GO term. ASP3, for example, is both a structural component of the cell wall and a metabolic gene, but is only annotated as the latter.

**Figure S3.**
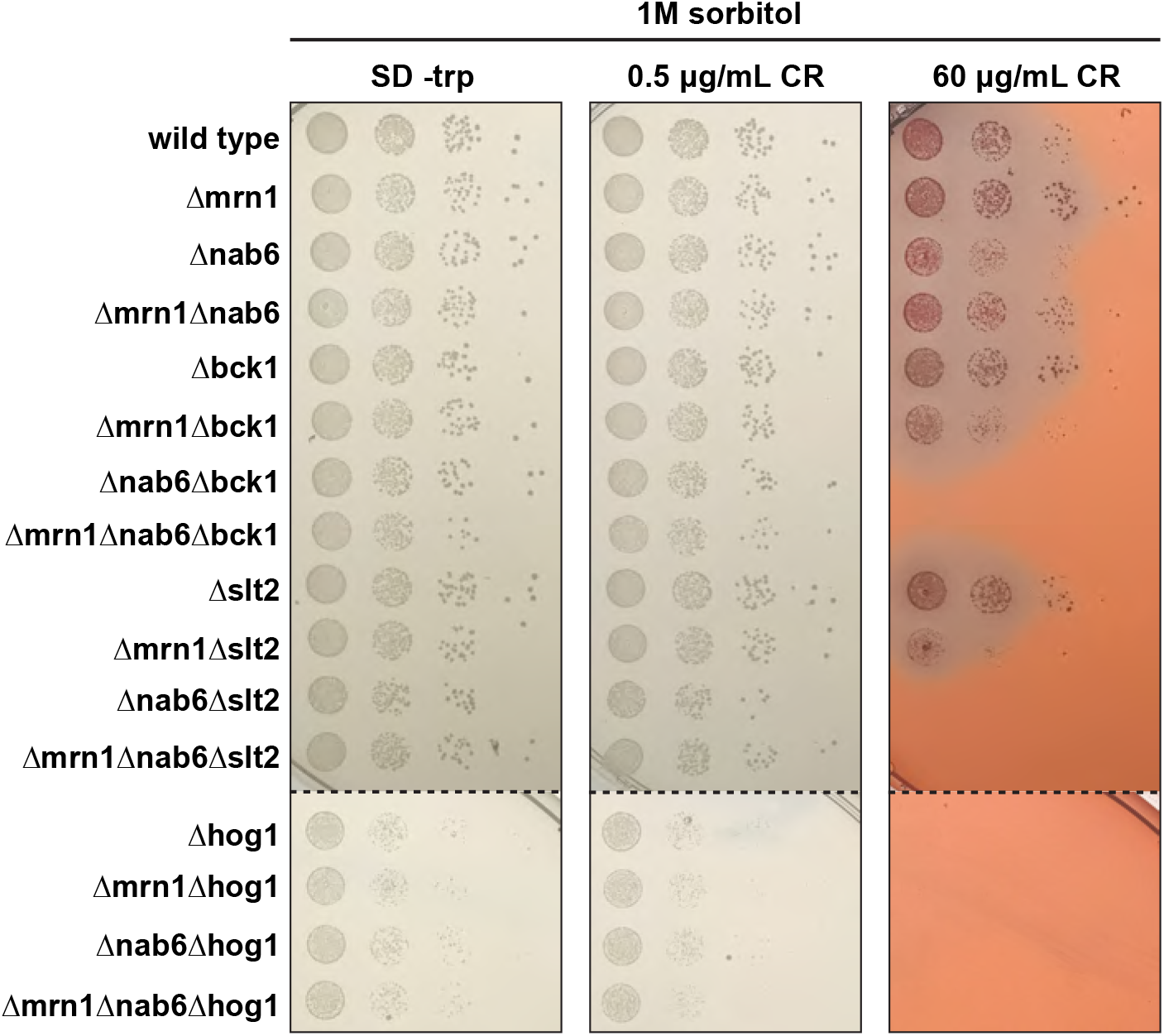
Related to Figure 2. Strains with combinations of gene deletions were tested for their ability to grow on plates with 0.5 μg/mL and 60 μg/mL Congo Red supplemented with 1 M sorbitol. Cells were grown for two days at 30°C.

**Figure S4.**
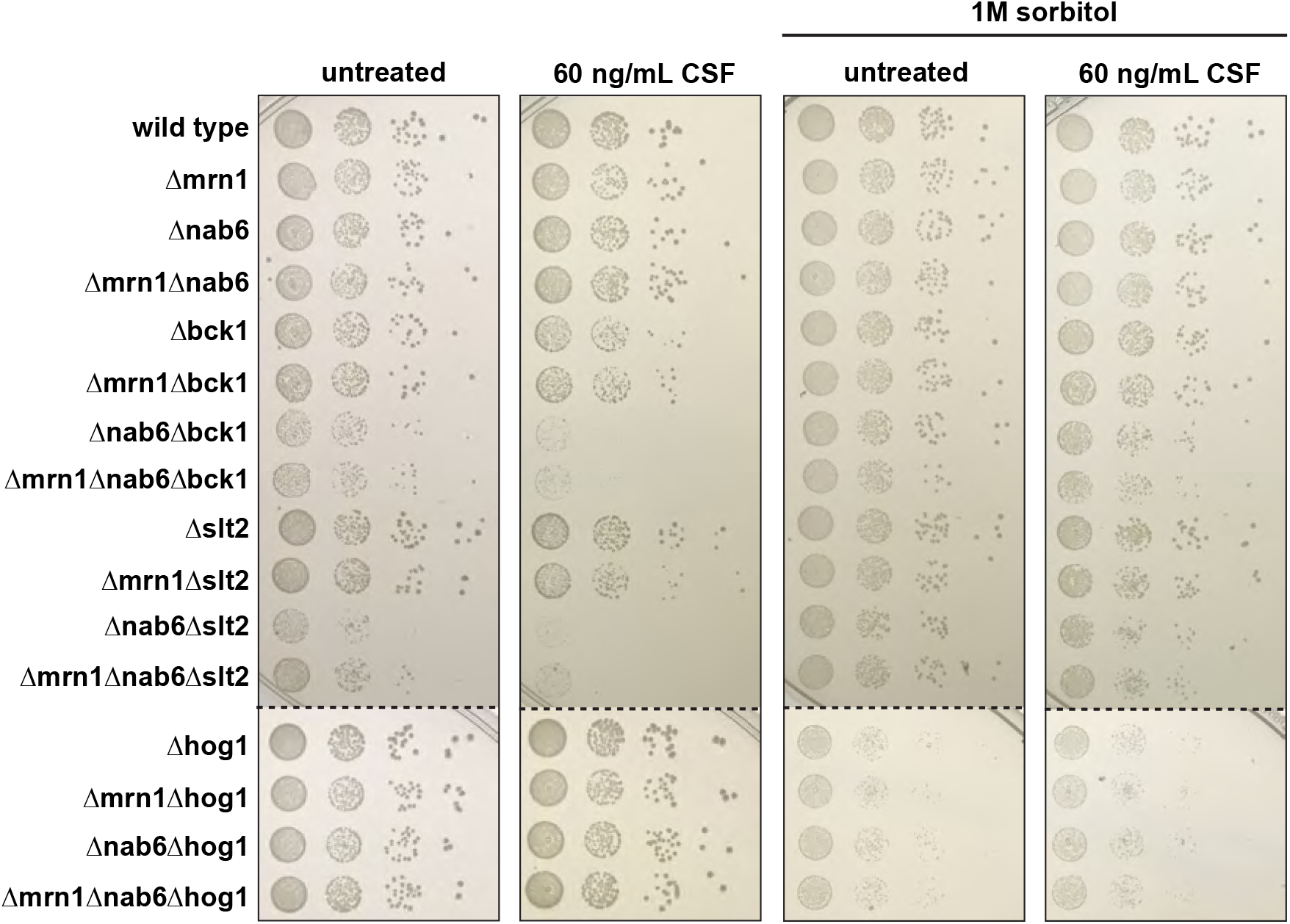
Related to Figure 2. Strains with combinations of gene deletions were tested for their ability to grow on plates supplemented with 60 ng/mL caspofungin (CSF) and with or without addition of 1 M sorbitol. Growth assays for cells on plates lacking CSF are included for comparison (identical to the images in Figs. 2B and S3). Cells were grown for two days at 30°C.

**Figure S5.**
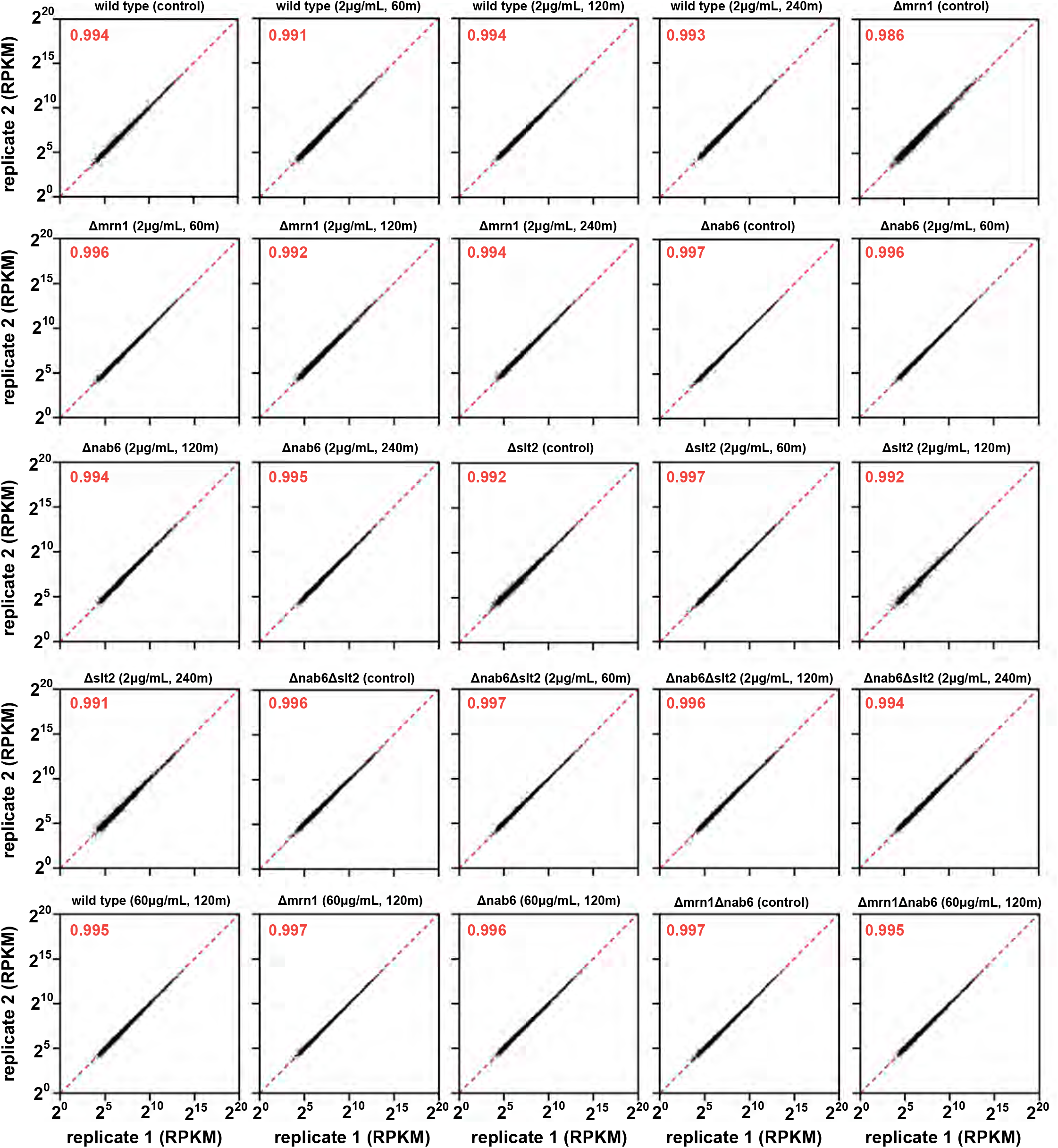
Related to Figure 3. Comparison of RNAseq replicate datasets for 5000 mRNAs. The red number indicates the spearman correlation coefficient.

**Figure S6.**
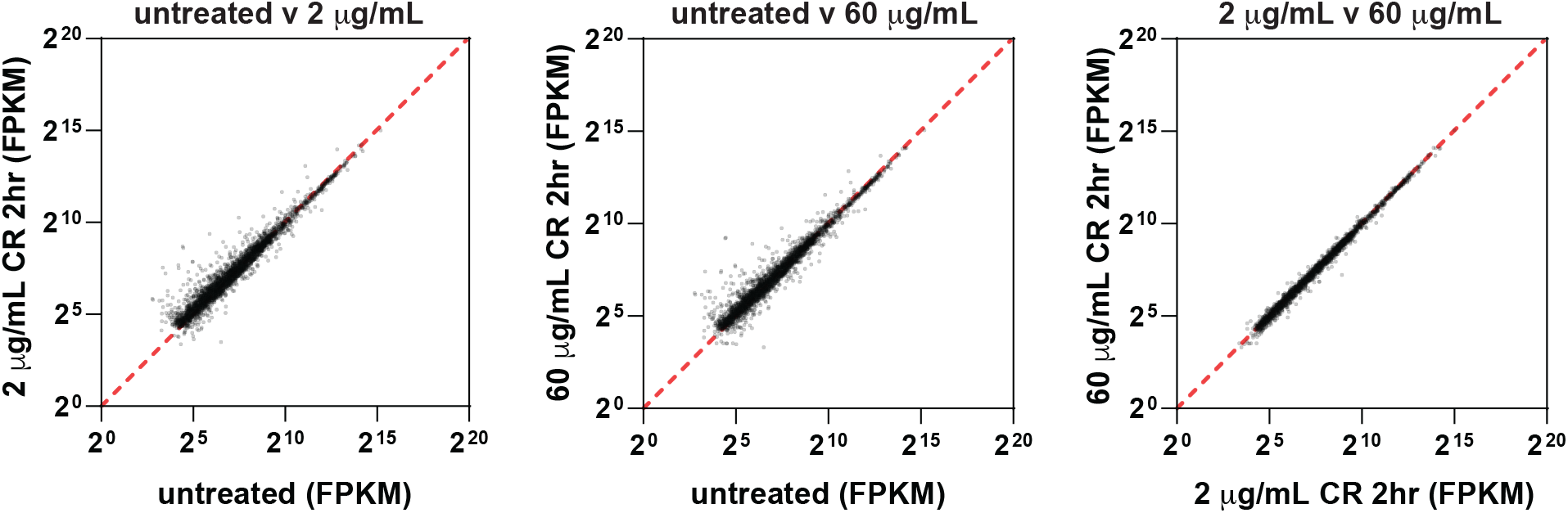
Related to Figure 3. Scatter plots showing comparisons between different RNAseq datasets for the wild type strain.

**Figure S7.**
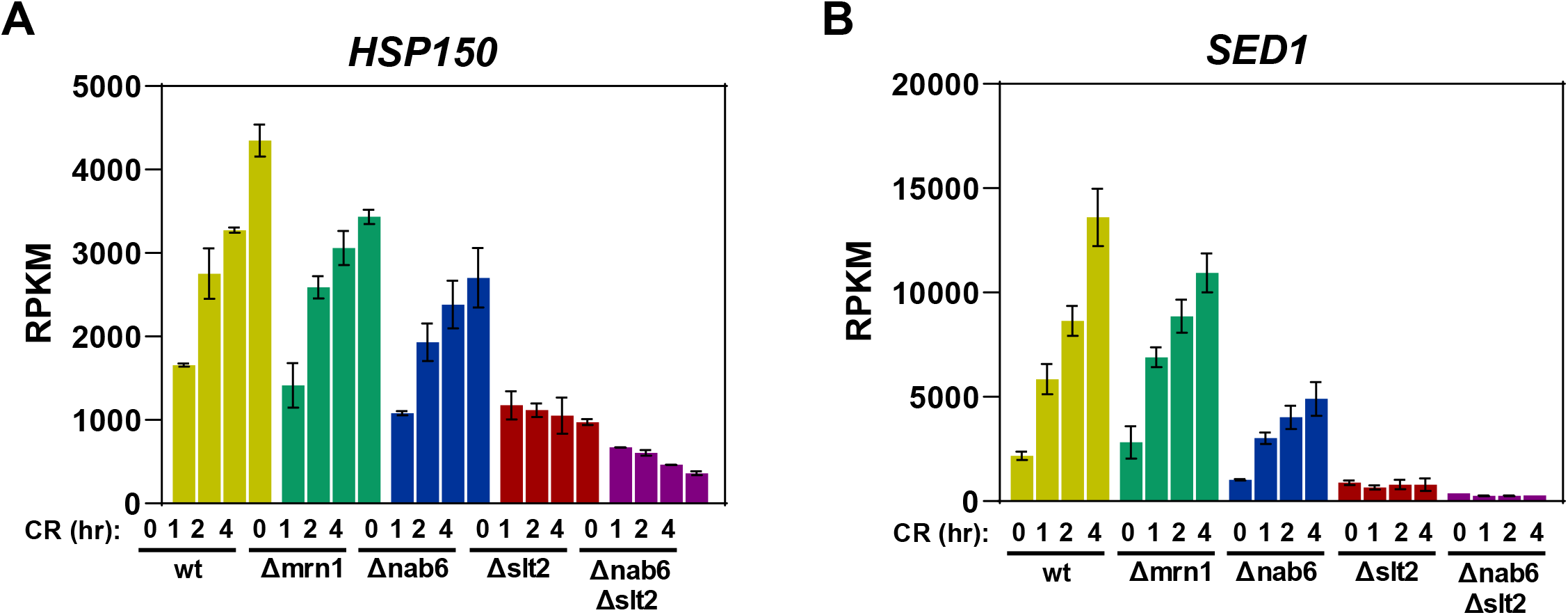
Related to Figure 3. (A-B) Bar graphs showing changes in (A) *HSP150* and (B) *SED1* mRNA abundance following CR treatment.

## Notes

### Competing Interest Statement

The authors have declared no competing interest.

